# Advances in Geometric Techniques for Analyzing Blebbing in Chemotaxing *Dictyostelium* Cells

**DOI:** 10.1101/526608

**Authors:** Zully Santiago, John Loustau, David Meretzky, Devarshi Rawal, Derrick Brazill

## Abstract

We present a technical platform that allows us to monitor and measure cortex and membrane dynamics during bleb-based chemotaxis. Using *D. discoideum* cells expressing LifeAct-GFP and crawling under agarose containing RITC-dextran, we were able to simultaneously visualize the actin cortex and the cell membrane throughout bleb formation. Using these images, we then applied edge detect to generate points on the cell boundary with coordinates in a coordinate plane. Then we fitted these points to a curve with known *x* and *y* coordinate functions. The result was to parameterize the cell outline. With the parameterization, we demonstrate how to compute data for geometric features such as cell area, bleb area and edge curvature. This allows us to collect vital data for the analysis of blebbing.

## 2 Introduction

Cells must modify their motile behavior when encountering varying conditions. They must travel through multiple environments as they participate in a variety of biological phenomena including foraging for food, embryogenesis, development, wound healing, immune response, and cancer metastasis. There are two distinct modes of motility cells utilize depending on their environment [1], [2]. When crawling on top of a substrate with limited resistance to movement, a two dimensional environment, cells use filopodia, lamellipodia, or pseudopodia as their main mode(s) of motility where actin is continuously cycled to the front of the cell, pushing the cell’s membrane forward in the direction of movement. When crawling through a substrate or between cells where resistance is higher, a three dimensional environment, cells use blebs as their main mode of motility. During bleb-based motility, the front of the cell makes a series of blister-like protrusions in the direction of movement where the cell’s membrane detaches from the actin cortex [3]. This is driven in part by the increased intracellular pressure associated with moving through a three dimensional environment. A variety of cell types have been shown to utilize bleb-based motility in three dimensional environments: skeletal muscle stem cells, zebrafish primordial germ cells, cancer cells, *Entamoeba histolytica* and *Dictyostelium discoideum* [4], [5], [6], [7], [8], [9] and [10].

The formation of a bleb follows three general steps with distinct membrane and cortex characteristics (Fig 1): 1) the membrane detaches from the cortex, making a blister-like protrusion at the cell front; 2) the new cortex begins forming at the new position of the membrane while the original cortex behind the detachment begins to disassemble; and 3) the original cortex vanishes where the new cortex is fully assembled and associated with the membrane.

**Fig 1.**
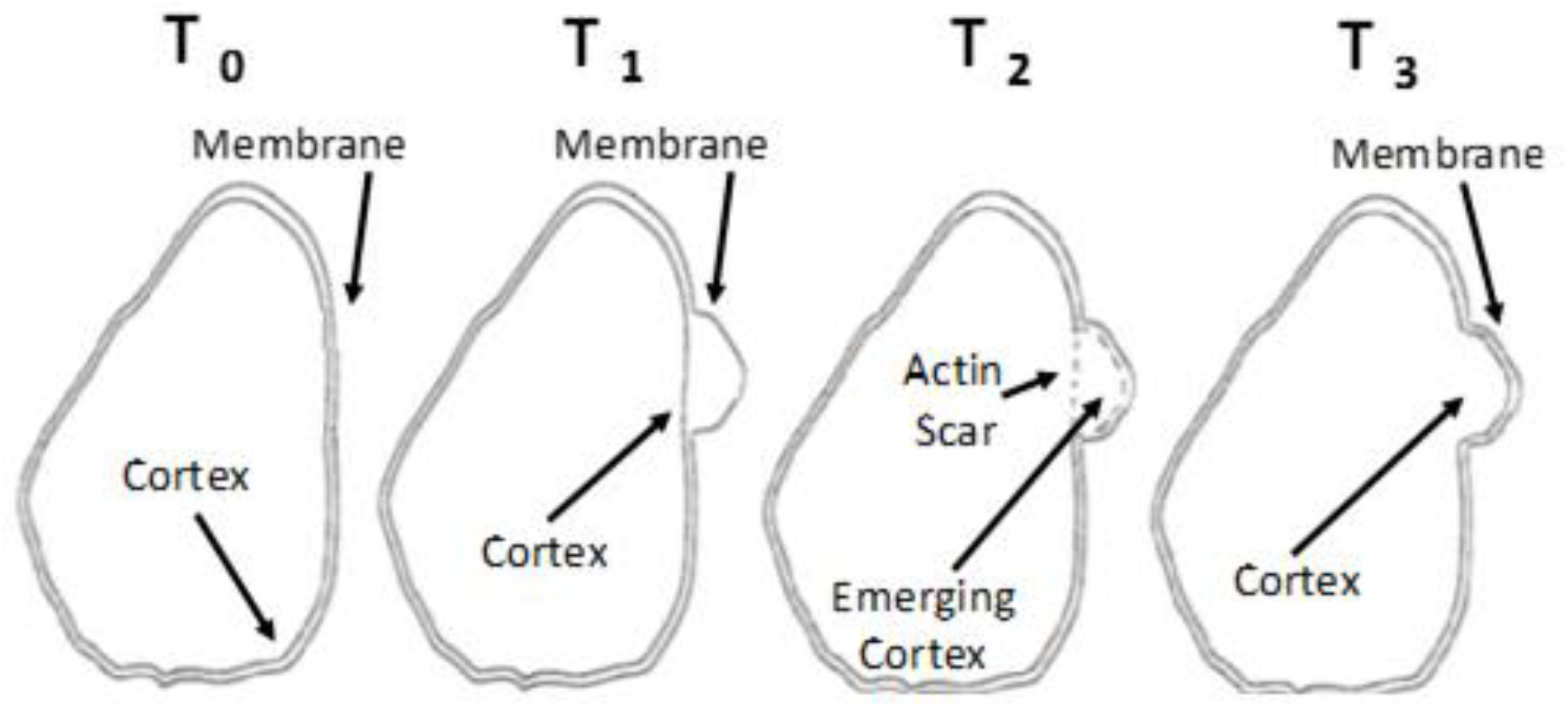
Bleb formation can be identified by cortex-to-membrane positioning. At *T*_1_, the membrane detaches from the cortex, initiating a bleb. At *T*_2_, an actin scar in the original location of the cortex disassembles as the cortex begins to reform at the new location of the membrane. At *T*_3_, the cortex is bound to the membrane in the bleb.

In recent years, several researchers have studied bleb formation from different points of view. In [11], Charras et al. collected biophysical data from the cell during blebbing, leading them to conclude that the blebbing process is the result of pressure changes. They proposed a model for bleb nucleation. However, these conditions were necessary but not sufficient. Using classical geometric constructs, Wooley et al. [12] modelled bleb development and were able to explain several bleb shapes that occur in nature. Guy and Strychalsky [13], [14] considered the same process from the point of view of computational fluid dynamics. By simulating the fluid flows, they were able to include secondary blebbing in their model. Zatulovskiy, Tyson, Bretschneider, and Kay [3] visualized the cortex and membrane dynamics during bleb formation in *D. discoideum* using an under agarose assay and introduced differential geometry via membrane/cortex curvature to the process. They showed that curvature does play a role as bleb location is biased toward areas of negative curvature. However, it is apparent from this work that there are other factors at play. Collier et al. [15] proposed that cell surface energy may help predict bleb nucleation sites, and that membrane curvature is only one factor in the cell surface energy calculations. This is in keeping with [16], where the authors correlate the presence of Myosin-II with bleb formation.

There are several proposed mechanisms necessary for bleb formation [17]. However, these do not fully explain directional bleb-based motility. Blebs appear as a consequence of the three dimensional environmental force transferred to increased hydrostatic force on the cortex/membrane complex. The central question is why the blebs appear on the anterior face leading to coordinated cell movement. Even if blebs tend to occur at sites of negative curvature [18] and [3] or high surface energy [15], these do not explain why blebs are more likely on the anterior face. Indeed, there are negative curvature segments and high energy locations throughout the cell boundary. In order to collect the data needed to address this conundrum, large numbers of blebbing cells need to be imaged at high enough resolution to visualize cortex and membrane structures. In addition, these images need to be analyzed to accurately measure and quantify membrane curvature and bleb size throughout bleb-based motility.

In this letter, we describe the technical procedures permitting us to collect images of the cortex and membrane during blebbing as well as the automated computer application that approximates the cell boundary with a B-spline and measures curvature, allowing for a high throughput geometric analysis of blebbing during chemotaxis. This letter is organized as follows. In the Materials and Methods, we state in turn, cell preparation, microscopy procedures, edge detection and geometric modeling functions. In the Results, we apply the methods to *D. discoideum*. In particular, we produce a parametrized representation of the cell boundary. We then proceed to show some applications of our methods. We conclude by tabulating blebs by edge curvature. Additionally, we include two Appendices in the Supplemental Materials. In Appendix A, we include videos of the cells we used for our study. In Appendix B, we include a mathematical result that justifies our choice of parameterization procedure. In the Discussion and Conclusion, we reflect on what has been done and the importance of this work.

## 3 Materials and Methods

### 3.1 Cell preparation

#### 3.1.1 Strain and culture conditions

All *D. discoideum* cells were grown axenically by shaking culture in HL5 nutrient medium with glucose (ForMedium) and supplemented with 100 IU/mL penicillin and 100 *μ*g/mL streptomycin (Amresco) at 150 rpm at 22°C. To visualize F-actin dynamics in wild-type cells, the Ax2 strain was used. Wild-type cells were transformed with the LifeAct-GFP plasmid, a generous gift from Dr. Chang-Hoon Choi, University of California. To make the construct [19], the LifeAct-GFP sequence [20] was amplified by PCR using the following primers:

> LifeAct – F, 5′ – AAAAGATCTAAAAAATGGGTGTCGCAGATTTG – 3′

and

> LifeAct – R2,5′ – TTTCTCGAGTTAAGCGCCTGTGCTATG – 3′.

The amplified LifeAct-GFP sequence was cloned into the BglII and XhoI sites of the EXP5(+) vector containing a neomycin-resistant cassette by electroporation as previously described [21]. Transformed cells were selected in HL5 media supplemented with 4 – 20*μ*g/mL G418 (Geneticin). Individual clones were isolated on GYP plates (0.2% peptone, 0.02% yeast extract, 2.2% agar, 0.1% dextrose, 19 mM Na_2_HPO_4_, and 30 mM KH_2_PO_4_) containing 4 μg/ml G418 on Br/1 bacteria. Clones were chosen for imaging based on similarity to F-actin LifeAct-GFP fluorescence labeling found in previous work in *D.discoideum* [22].

To make the cells competent for cAMP chemotaxis, 1 × 10^7^ log phase Ax2 LifeAct-GFP expressing cells were collected and washed in PBM (20 mM KH_2_PO_4_, 10*μ*M CaCl_2_, 1 mM MgCl_2_, pH 6.1, with KOH) three times as previously described [23]. The cells were resuspended in 1 mL PBM and plated at 22°C on a 0.8*μ*m pore size, 47mm in diameter nitrocellulose filter pad (Millipore) on top of two 6*μ*m pore size, 125mm in diameter Whitman paper filter pads (GE HealthCare) that were pre-moistened with PBM. After 5 to 6 hrs, the starving cells were removed and suspended in 1mL PBM.

#### 3.1.2 cAMP under agarose assays

Two types of cAMP under agarose assays were used: 1) to visualize membrane and cortex positions and 2) to visualize bleb progression after nucleation.

To visualize membrane and cortex positions in bleb-based motility in response to cAMP chemotaxis, an under agarose assay developed by Zatulovskiy, Tyson, Bretschneider, and Kay [3] was used with the following modifications. A preheated (90° C, 1 min) number 1 German borosilicate sterile 2 well chambered coverglass slide (ThermoFisher) was loaded with 750*μ*L of melted 0.5% or 0.7%Omnipur^®^ agarose (EMD Millipore) laced with 1mg/mL of 70,000 MW Rhodamine B isothiocyanate-Dextran (Sigma-Aldrich). Once solidified, half of the gel was removed from the well and the remaining portion was slid across to the middle of the chamber, creating two wells with one on each side of the gel. To create the cAMP gradient, 4*μ*M cAMP was loaded into one well and incubated for 40 minutes. 1 × 10^5^ to 2 × 10^5^ cells competent for cAMP chemotaxis were loaded into the other well and allowed to settle and crawl under the agarose for 30 minutes prior to imaging.

To visualize bleb progress after nucleation, the same cAMP under agarose assay to visualize membrane and cortex positions was used; however, no RITC-Dextran was added to the agarose gel.

### 3.2 Imaging and microscopy

#### 3.2.1 Live imaging and microscopy

All imaging data were acquired using a Leica DMI-4000B inverted microscope (Leica Microsystems Inc.) mounted on a TMC isolation platform (Technical Manufacturing Corporation) with a Yokogawa CSU 10 spinning disc head and Hamamatsu C9100-13 ImagEM EMCCD camera (Perkin Elmer) with diode lasers of 491 nm, 561nm, and 638 nm (Spectra Services Inc.) [24]. LifeAct-GFP and RITC-Dextran were excited using the 491nm and 561nm lasers, respectively. Images were taken over the course of 30 seconds using 80x magnification (40x/1.25–0.75 oil objective with a 2x C-mount) or a 100x/1.44 oil immersion objective at maximum camera speed with exposure times of 0.800 seconds for GFP and 0.122 seconds for RITC channels, resulting in intervals of 1.66 seconds for the visualizing membrane-to-cortex position assays and intervals of 0.800 seconds for the bleb progression assays as only the GFP channel was needed. Images were acquired using Volocity 5.3.3 (Perkin-Elmer) and processed in ImageJ only for intensity plot analysis.

#### 3.2.2 Intensity plots to determine cortex and membrane positions

ImageJ signal intensity plots were used to confirm positions of the membrane and cortex in all steps in bleb formation. Frames acquired from both channels were put together in a single stack where a line was drawn across the middle of the putative bleb. The line was drawn starting from the cytoplasm just prior to the cortex and extending past the bleb in question. From ImageJ, the Plot Profile feature (Analyze > Plot Profile) was used to generate an intensity plot for each frame in the stack.

### 3.3 Data conversion - Microscopy to Cartesian coordinates

Data conversion was implemented using *Mathematica* standard library, image processing functions. The goal was to convert digital microscopy output into Cartesian coordinates.

#### 3.3.1 *Mathematica* image processing functions

The *Mathematica* functions we use for image processing are as listed.

- *ImageTake* was used to clip the image to eliminate extraneous neighboring objects. The parameter setting depended on the location of the desired image in the field of view.
- *FillingTransform* was used to remove small isolated black areas in the field of view. This function blended these with neighboring gray areas to form a more uniformly gray area.
- *ImageAdjust* scaled the brightness of the image to the unit interval [0, 1].
- *Blur* is a standard computer graphics procedure that modified pixel values by interpolating values at neighboring pixels.
- *Binarize* created a bi-chromatic, black background, white central feature. The threshold parameter used to distinguish between gray levels was set in the range 0.09 to 0.21. The choice of parameter depended on the sharpness of the microscopy output. For sharp images, we used values near 0.1. For fuzzy images, values near 0.17 to 0.21 removed the spurious output.
- *DeleteSmallCompnents* corrected for anomalies in the Binarize output, small connected components of white within the black or vice-versa. The connected components were identified using cluster variance maximization or Otsu’s algorithm.
- *EdgeDectect* identified the edge of interest from the bi-chromatric image. This function implemented the Canny algorithm.
- *PixelValuePositions* produced the cell boundary in Cartesian coordinates.
- *FindShortestTour* transformed the list produced by PixelValuePositions into an ordered circuit.

### 3.4 Geometric functions

#### 3.4.1 Parameterizing the cell boundary

We transition now to the mathematics. Given a planar object and set of points on the boundary, *B_i_*, *i* = 1, …, *n*, the procedure arrives at a parameterized representation of the object. All other geometric functions will be built from this representation of the object.

A cubic B-spline is determined by guide points, *B_i_*, *i* = 1, …, *n* and cubic basis functions,

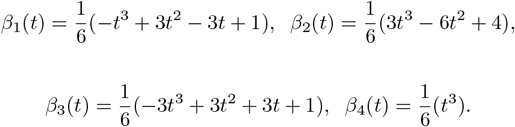

The resulting interpolating curve is given in segments. Each segment is determined by four of the guide points. The equation for the *i^th^* segment uses guide points *B_i_*, *B*_*i*+1_, *B*_*i*+2_, *B*_*i*+3_.

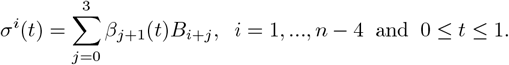

If the points lie on a curve, then the B-spline is a parameterized approximation of the curve.

This representation has the advantage that it has a formula. A curve with a known formula is referred to as a *parameterized* curve. For the case of the B-spline, it is a polynomial in each output variable. We can differentiate it, integrate under it. We can compute the length of any part of the curve, compute the tangent vectors and the curvature at locations. We can identify a sequence of equally spaced points on the curve.

Furthermore, we used Delaunay triangulation to compute the area inside a parameterized curve. *Delaunay triangulation* was implemented in *Mathematica*. By triangulation of a planar region, we mean the region divided into disjoint triangles. When given a point sequence on the curve, this function returned the triangulation of the convex hull enclosed by the points. This can subsequently be modified to return the area inside the curve.

There is a *Mathematica* routine to do B-splines. However, we preferred to implement the equation for B-spline directly. This gave us more control over the output.

## 4 Results

### 4.1 Image acquisition and bleb identification

#### 4.1.1 Cortex and membrane positions were visualized using the under agarose assay

In order to visualize cortex and membrane dynamics of a chemotaxing cell, we modified an under agarose assay developed by Zatulovskiy, Tyson, Bretschneider, and Kay (2014) [3]. Briefly, starved cells were filmed as they crawled under a slab of agarose towards a source of cAMP. The cortex was visualized by labeling with LifeAct-GFP and the membrane was visualized by the shadow created by the cell against the agarose slab that contained RITC-dextran. Fig 2 shows a single time point of GFP and RITC channel images of a wild-type cell chemotaxing under a 0.5%agarose gel. LifeAct-GFP labeled the cortex as well as other F-actin structures (Fig 2, left). The cell produced a negative against the gel where the cell blocked the RITC signal. This provided an indication of the edge of the membrane where the negative ceased and the RITC signal intensified (Fig 2, center). Merging the GFP and RITC images provided cortex and membrane positions, which allowed the identification of a putative bleb as a blister-like protrusion mostly free of actin was observed (Fig 2, right). The full video of this cell can be found in Appendix A (Video 1).

**Fig 2.**
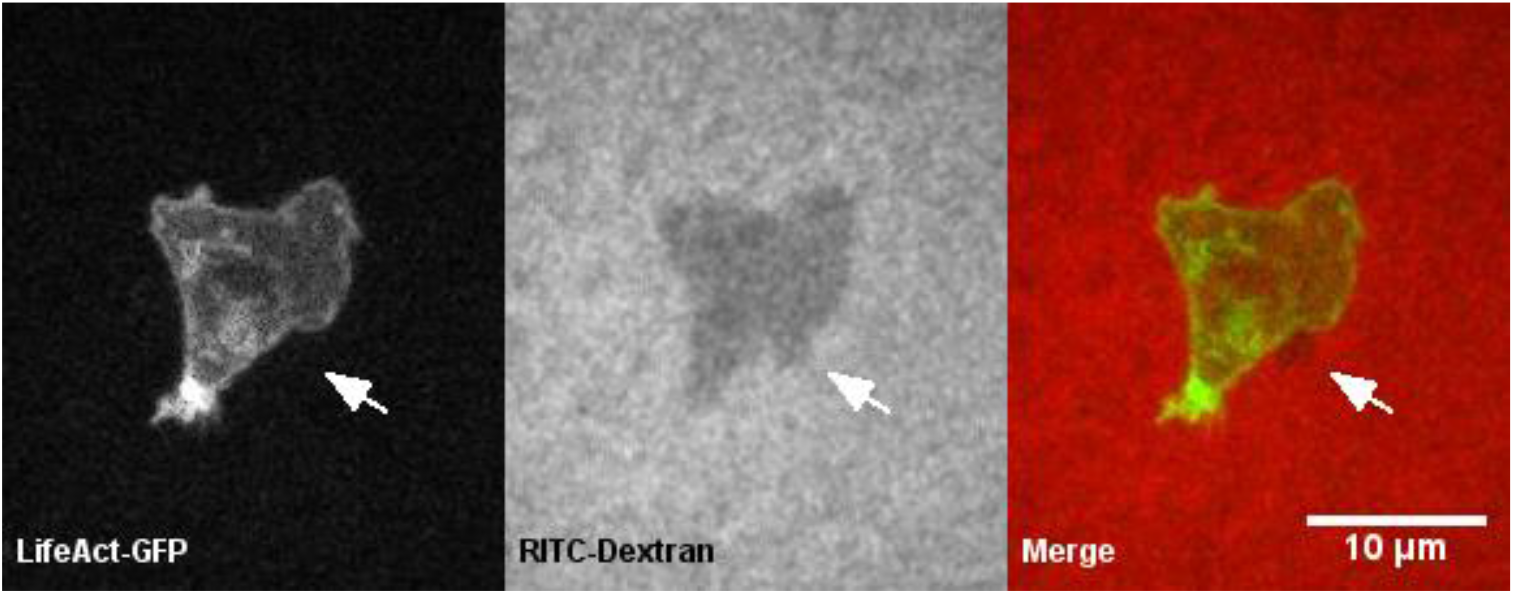
The under agarose assay simultaneously visualized cortex-to-membrane positions of a putative bleb. A single time point of a wild-type cell crawling under 0.5 percent agarose gel is shown. Left, LifeAct-GFP labeled the cortex’s position as well as other actin structures. Center, the RITC-Dextran in the gel was blocked by the cell, providing the cell’s edge and, therefore, membrane position. Right, a merge of the two channels provided cortex-to-membrane position where LifeAct-GFP and RITC-Dextran were false colored green and red, respectively. A putative bleb lacking actin structures was observed observed (white arrow).

#### 4.1.2 ImageJ intensity plots confirm cortex and membrane positions

In order to verify that the membrane detached from the cortex, forming a bleb, the membrane and cortex positions must be identified quantitatively. To accomplish this, we used ImageJ intensity plots (Fig 3). Using the same time point for each channel shown in Fig 2, we simultaneously determined the positions of both the membrane and cortex quantitatively (Fig 3). The signals of the LifeAct-GFP (dashed line) and RITC (solid line) were quantified along a drawn line (white) on the images. The cortex’s position was identified by the highest GFP signal intensity peak, which occurs at approximately 1.1 microns. The membrane’s position was identified by the negative created by the cell where the edge of the cell was represented by the sharp increase in RITC signal intensity from the cell to the gel. For the membrane, a drastic increase in signal intensity was seen at approximately 2.9 microns. The merge shows the location of the membrane to the cortex as indicated by the distance, approximately 1.8 microns, between the GFP and RITC peak intensities (Fig 3).

We then questioned whether we could use this method to determine cortex-to-membrane positions during bleb formation.

**Fig 3.**
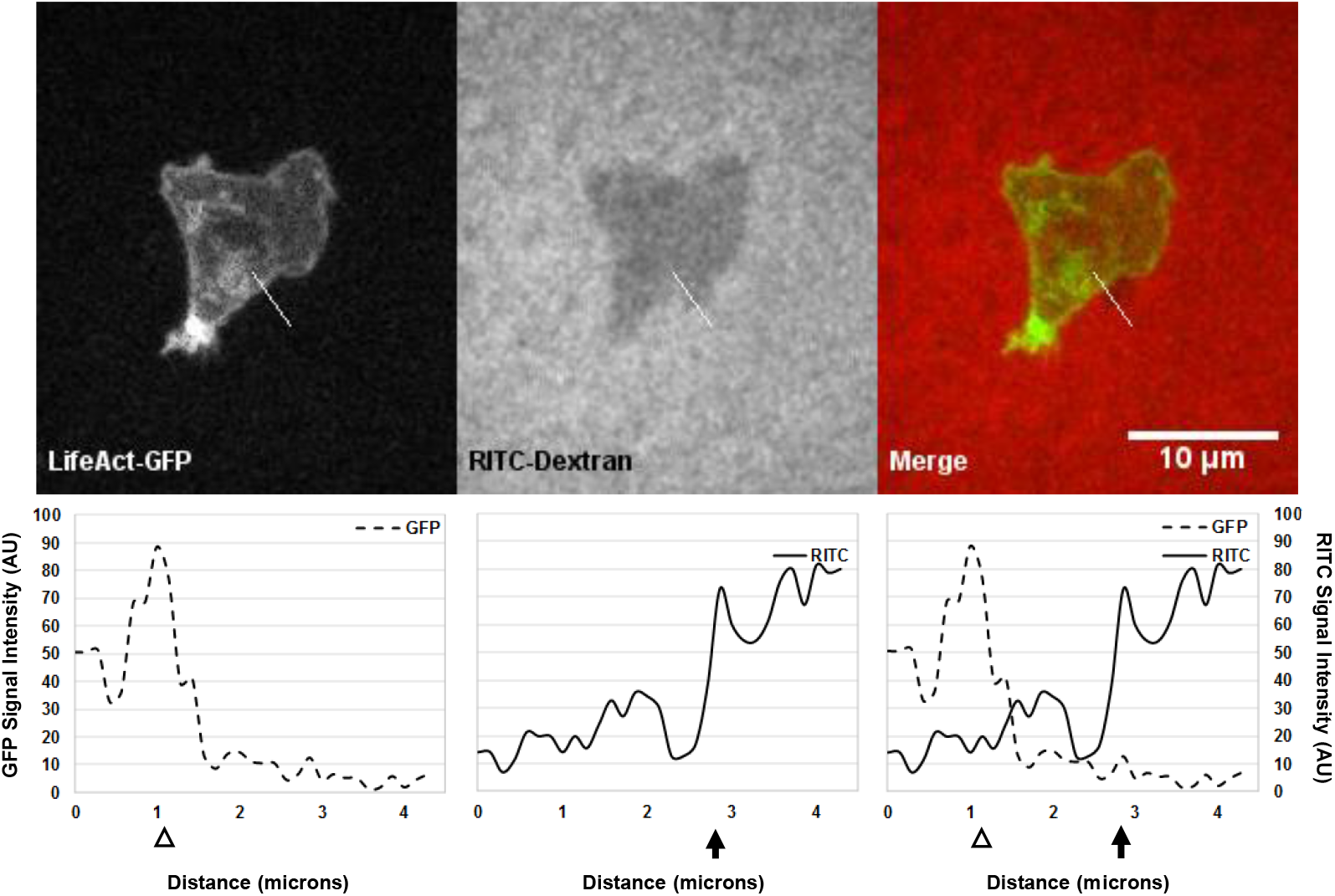
ImageJ intensity plots quantitatively confirm membrane and cortex positions of a putative bleb. A single time point of a wild-type cell crawling under 0.5 percent agarose gel from Fig 2 with respective intensity plots of the signals along the drawn line (white line) for each channel is shown. Left, the LifeAct-GFP signal (dashed line) indicated the cortex’s position (white arrowhead) at 1.1 microns. Center, the RITC-Dextran signal (solid line) indicated the membrane’s position (black arrow) at 2.9 microns where the cell’s membrane ends and the agarose begins. Right, the merge of the two channels shows membrane detachment from the cortex by a distance of 1.8 microns.

#### 4.1.3 Intensity plots confirm bleb formation dynamics

In order to confirm the formation of a bleb by the cell shown in Fig 2, we determined membrane and cortex positions over the course of several seconds encompassing before and after the putative bleb formation and compared that to the membrane and cortexes’ expected dynamics for bleb formation and completion (Fig 1, [10]). As shown in Fig 4 at *T*_0_, the cortex and membrane were in close proximity at 1.0 and 1.3 microns, respectively. At *T*_1_, the membrane’s sharp increase in RITC intensity shifted to 2.9 microns while the cortex peak remained in its original location; thus the membrane detached from the cortex, indicating the start of a bleb. At *T*_2_, the membrane’s sharp increase in RITC intensity shifted slightly further to 3.3 microns where the original cortex’s peak signal lessened, representing the actin scar characteristic, and another GFP-peak at 3.1 microns was seen in close proximity to the shifted membrane, indicating that a new cortex was being assembled in close proximity to the new membrane location. At *T*_3_, the shifted membrane moved a little further to 3.4 microns whereas the new cortex at 3.1 microns was fully assembled, and the actin scar lost its signal intensity, demonstrating that the original cortex completely dissipated. As shown, all characteristics of bleb formation were unambiguously identified by this method. To further support this method, a non-blebbing region on the same cell shows the cortex/membrane complex being maintained over the same time course (SFig. 2). The full video of the cell can be found in Appendix A, Video 1.

**Fig 4.**
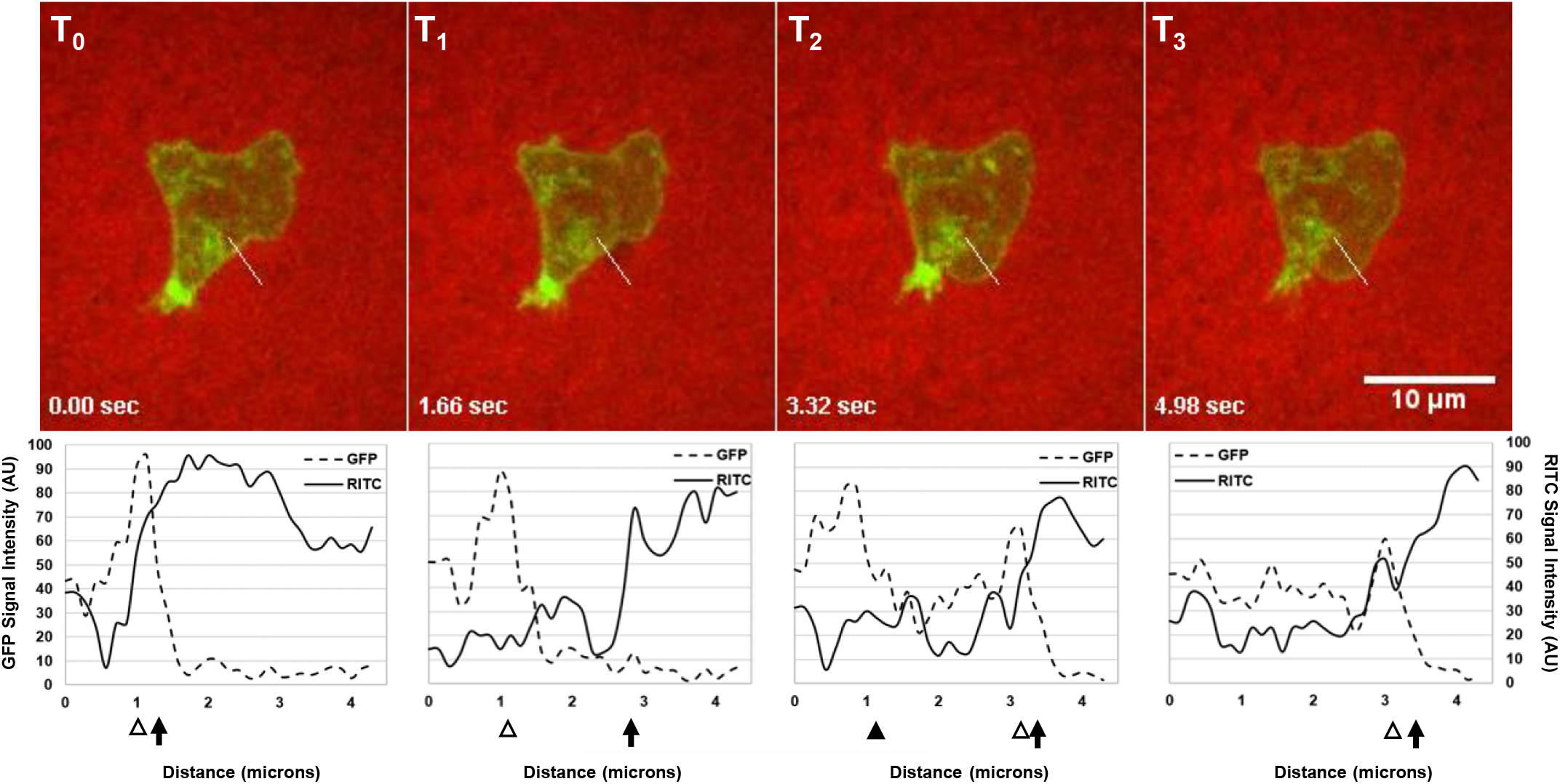
ImageJ Intensity plots confirmed cortex-to-membrane positions throughout bleb formation of a putative bleb. Merges of GFP and RITC time points representative of steps in bleb formation (*T*_0_-*T*_3_) with their respective intensity plots (dashed and solid lines) from the same cell in Fig 2 confirmed bleb formation quantitatively in real-time along the line drawn (white). Cortex positions (white arrowhead), the actin scar characteristic (black arrowhead), and membrane (black arrow) positions are indicated.

### 4.2 Image processing

#### 4.2.1 General statements

A *cell image* refers to a sequence of microscopy images taken every 1.66 seconds for the cortex and membrane position assays and 0.800 seconds for the bleb progression assays. These images are still frames from a digital video of the cell. It is important to note that for each cell, we process stills or frames identically in all aspects. Videos of all cells presented in this letter can be found in Appendix A.

In Image Processing, we take this sequence and identify the boundary points for each still or frame. In particular, we set up a coordinate system and then identify the coordinates for the boundary points. The point sequences produced in this step are subsequently input into the geometric modeling component described in the next section.

A note on what is being detected in our analysis. It is apparent we are locating the cell cortex in the GFP images. Since the membrane and cortex are usually tightly joined, in most cases we are locating the cell *boundary*, the membrane/cortex complex. The only exception occurs during a bebbing event. In this case, the membrane extends without any apparent supporting cortex. Generally, it takes about a second (the time to the next still) for the cortex to form at the extended membrane. Even before then, actin debris is often visibly present in the new bleb. Our process identifies this material. The result is often strangely shaped. We have learned to recognize this as a bleb in progress. (See SFig 3 through SFig 5.)

The following computer functions when executed sequentially describe an automated process that renders the microscopy image as a parametrized curve. (See Section 4.3 Geometric modeling.)

#### 4.2.2 Steps Toward Digitization

Images were first cropped using the *ImageTake* function. The purpose of this step was to remove extraneous nearby objects. The resolution and magnification of the images were not altered during this step. See Fig 5-1 and Fig 5-2. Of course all images in the sequence were cropped identically.

**Fig 5.**
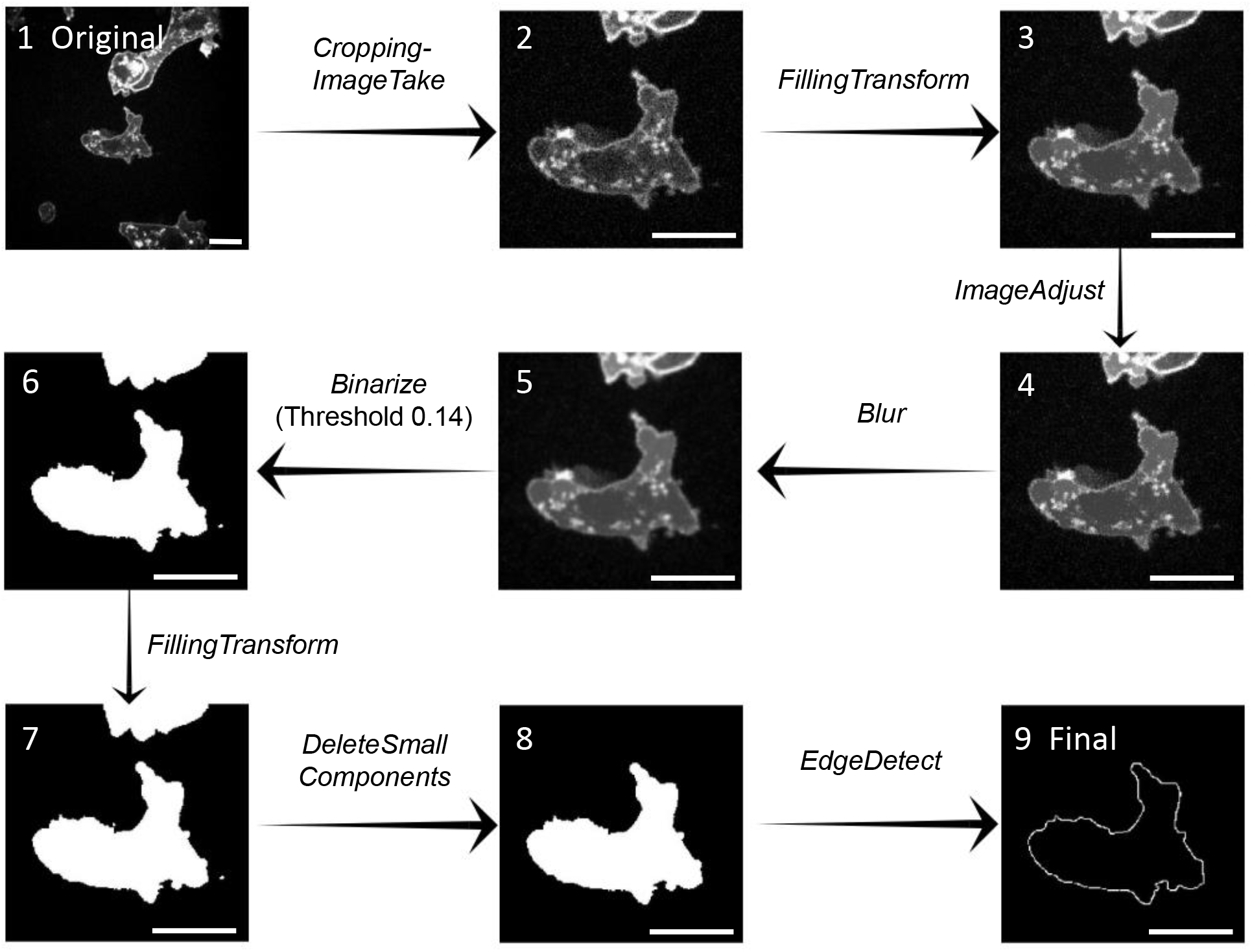
Image Processing: Microscopy → Digitization Flowchart. 1. The original microscopy output. 2. The cropped image. Only unnecessary background was removed. 3. The result of *FillingTransform*. Very small dark features were blended with adjacent areas to form a more uniformly gray cell interior. 4. *ImageAdjust* rescaled the brightness component within the unit interval, [0, 1]. 5. The blurring operation modified areas where high intensity and low intensity were adjacent. 6. *Binerize* reset the image pixels into 0 or 1. The *Binerize* parameter set the threshold determining which pixels were set to white (1) and black (0). 7. The filling transform was reapplied. 8. *DeleteSmallComponents* removed any unwanted artifacts, white areas inside the black and black inside the white. This ensured there is only one edge to detect. 9. *EdgeDetect* identified the edge between the black and white regions. Scale bars identify 10 micron lengths.

The next goal was to prepare the image for the edge detection step. We wanted to produce a bi-chromatic image where the cell was white and the external matrix was black. To this end, we eliminated unwanted detail or reduced noise until the only remaining detail was the cell edge.

In our setting, LifeAct-GFP was expressed in the cell to illuminate the F-actin. During image acquisition, signals produced by the labeling were visualized by using gray-scale where the highest signal of the label was white. The result was a very bright, almost white, actin cortex at the cell boundary. The interior of the cell with only isolated actin polymers was mostly a dull gray punctuated with bright specks. We implemented *FillingTransform*. The result was to reset the isolated dark spots to match the surrounding region. Hence, the cell interior was more uniform and more easily distinguished from the cortex. Compare Fig 5-2 and 5-3.

We used *ImageAdjust* to rescale the brightness to values in the unit interval [0, 1]. There was little visual change, see Fig 5-4. This is a useful technical modification in preparation to the following steps.

The final noise reduction step employed a blurring transformation, *Blur*. Compare Fig 5-4 and Fig 5-5. This transformation diminished the contrast within the cell by interpolating brightness values at each pixel using a weighted average of the surrounding pixels. This reduced isolated actin fragments.

We begin edge detection by applying *Binarize*. The *Mathematica* function takes the bi chromatic image and assigns a value of zero and one to each pixel. The assignment depends on the brightness level as determined by a user set parameter. The choice of parameter is governed by the the sharpness of the image. The result was a white cell region and black background. *Binarize* output is illustrated in Fig 5-6. The same parameter, once selected, was used for the entire cell sequence.

At this point, we are nearly in position to do the edge detection. We removed anomalous artifacts that could confuse the edge detection. We did this in two steps. First we re-executed the filling transform followed by *DeleteSmallComponents*. This last application identified and removed small connected white regions in the black background or vice-versa. See Fig 5-7 and 5-8.

These previous two steps are often unnecessary. For our illustration, we have chosen a case where *Binerize* output leaves several disconnected white regions. In this case, *DeleteSmallComponents* removed the anomalies.

We are now ready to do the edge digitization. The function *EdgeDetect* identifies the points at the transition between the black and white regions. See Fig 5-9.

The final two steps are not included in the flow chart as they do not produce visual change. *PixelValuePosition* introduces the coordinate axis and assigns *x* and *y* coordinates to the physical boundary point set. The resulting set was not ordered as a circuit around the cell. Rather the points are ordered lexicographically for values of (*x*, *y*).

We sort the points identified by *PixelValuePositions* to form a boundary circuit using *FindShortestTour*. Often the cell shape is too complex and the *FindShortestTour* fails. We remedy this by separating the boundary point set, applying the function separately to each segment and then concatenating each piece to the whole.

### 4.3 Geometric modeling

#### 4.3.1 B-Spline fit

Our goal is to parameterize the cell boundary. Of course that is impossible. Rather, we approximate the cell boundary with a parameterized curve.

We begin by determining that the cell boundary is suitable for approximation by a B-spline. For this purpose, we identify two necessary assumptions. These statements will connect the mathematics to the biological observation. We will then introduce the B-spline parameterization and argue that it does (in a mathematical sense) parameterize the cell shape. Note that Lipschitz continuity is defined in Appendix B. Some of this material is necessarily technical.

*Assumption 1: The points returned by EdgeDetect actually lie on the cell boundary*.

*Assumption 2: The curve formed from the cell boundary is locally Lipschitz continuous*.

We begin our discussion of Assumption 1 by considering the correctness of the *EdgeDetect* procedure.

The correctness of *EdgeDetect* depends on the *Binarize* function. It assigns a value of zero and one depending on a user defined parameter which is determined using visual cues. The value 1 is assigned at and inside the cell boundary. Obviously, the parameter setting is sometimes flawed. Since we repeat the same parameter value for each still or frame associated to the cell image, then the same error is repeated at each still for the same cell. We expect that in a study using a large number of cells, the error will manifest as random variations with mean zero. Hence, the same will be true for the B-spline based on these points. For this reason, we may suppose that the *EdgeDetect* function is correct and adopt Assumption 1 as necessary support for the mathematics.

Regarding Assumption 2, we note that the cell boundary is smooth (differentiable). Indeed, the edge is formed by the membrane/cortex complex and shaped by local internal pressure [11]. As such, it cannot form a singularity. Even at the end of filopodium, the edge seems to round over the actin extension. In particular, we do not expect a cusp on the *D. discoideum* edge. This leads to the assertion that each point on the boundary lies on the graph of a function *f*(*x*) = *y* or *g*(*y*) = *x*, and this can be done so that the derivatives satisfy ║*f*′║ ≤ *D* and ║*g*′║ ′ *D* for some fixed constant *D*. This condition is necessary but not sufficient for local Lipschitz continuity. It is, however, close enough to lead us to accept Assumption 2.

We proceed to fit a cubic B-spline to the sequence of boundary points. The B-spline diverges from the actual boundary no more than the error bound, *K h* as computed in Appendix B. Here *K* is a constant and *h* is the mesh parameter, the maximum distance between successive guide points.

We want a B-Spline because of the smoothness properties. In particular, a cubic B-spline is twice continuously differentiable. Hence, it is rectifiable (has well defined length) and has well defined curvature everywhere. Furthermore, the error bound allows us to conclude that as *h* goes to zero, the distance between the curve and the boundary sequence goes to zero. This statement provides justification for the conclusion that the B-spline parameterizes the cell boundary. In Fig 6A, we compare the microscopy to the EdgeDetect to the B-spline versions of a cell.

**Fig 6.**
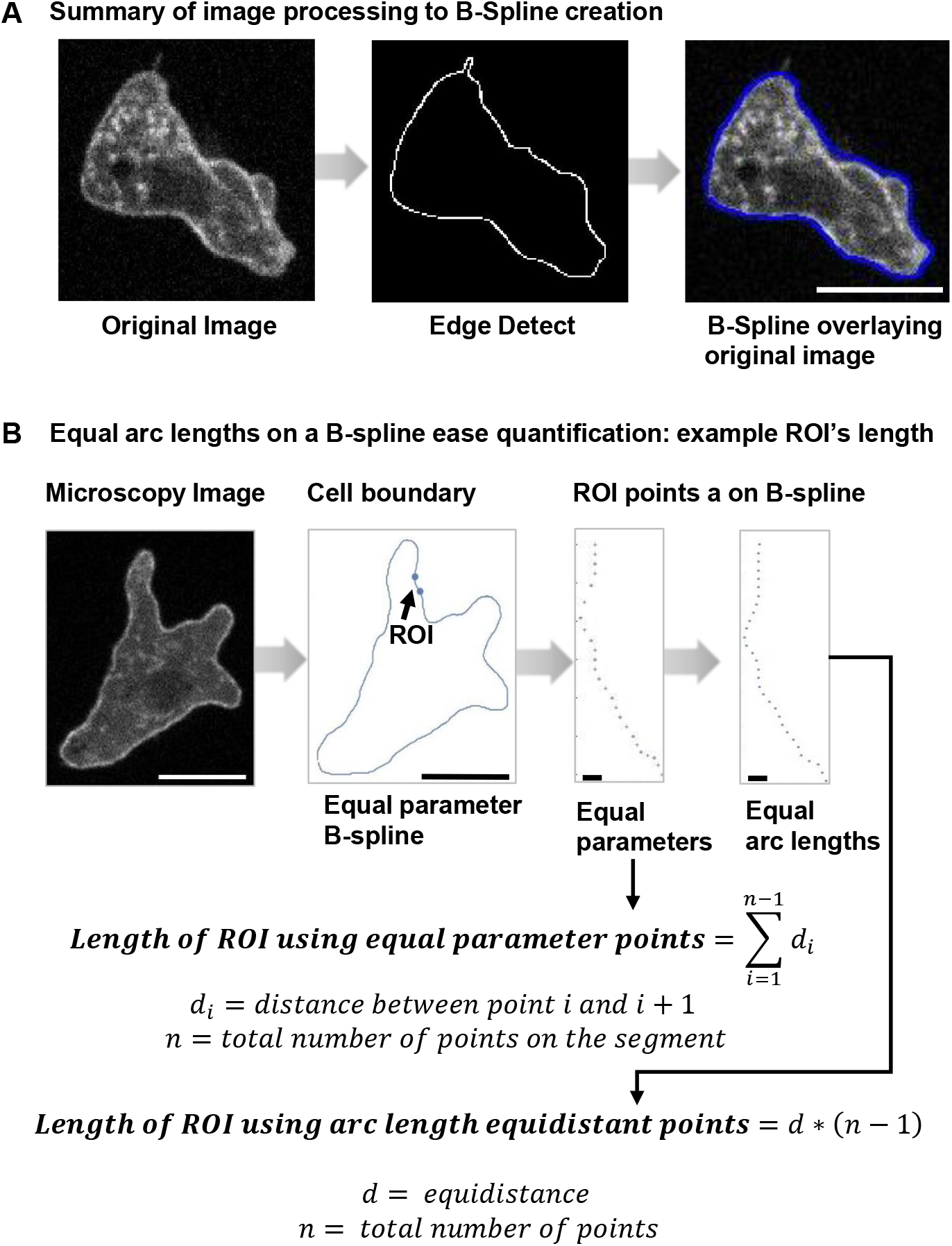
Creation of B-Splines and Equal Arc Length Points. (A) Three views of a cell: the microscopy image, the edge detect image, the B-spline plot overlaying the microscopy image. (B) The entire cell followed by two views of the boundary segment. The ROI view shows points on the B-spline at equal parameter increments, *t* = 0.5, and the same segment with points at equal arc length intervals, *δ* = 0.5 pixel units and *ϵ* = 0.001 pixel units. Large scale bars identify 10 micron lengths; small scale bars identify 1 micron.

#### 4.3.2 Equal spaced points

With parameterized representation of the cell boundary by a rectifiable curve, we are able to determine a sequence of equally spaced points on the cell edge. Points on the B-spline at equal parameter increments are not equally spaced. An equally spaced sequence can be used to regularly distribute a marker or to compute finite differences. In [3] such a sequence of boundary points was necessary for a membrane energy function.

Given a point on the B-spline, we select the next point along the curve at a preset distance *δ* and a maximum error *ϵ*. The selection is made using a binary search. In particular, given a point *P_i_*, we test points on the B-spline until we find one, *P*_*i*+1_ so that the distance between *P_i_* and *P*_*i*+1_ is between *δ* ± *∊*. Then we continue the process until we have completed the circuit.

The algorithm is complicated as the cubic B-spline has segments. We accommodate for the B-spline segments by translating the residual distance at the segment end forward to the next segment.

In Fig 6B, we see two views of a cell boundary segment. In once case, we have points at equal parameter spacing, in the other we have points at equal arc length. For instance, it is easier to compute distances along the cell boundary using the sequence of equally spaced points.

In the following subsections we illustrate the usefulness of the B-spline representation for calculating cell area, bleb area and relative curvature. In each case, there is an ongoing research project in our lab the uses these geometric procedures.

#### 4.3.3 Calculating area

We compute area by triangulating a region, dividing it into triangles and then summing the area of the individual triangles. If the region has a parameterized boundary, then we apply Delaunay triangulation to a sequence of boundary points. For instance, a sequence of regularly spaced boundary points. The Delaunay function in *Mathematica* returns the triangulation for the convex hull of the region defined by the points. See Fig 7A center.

**Fig 7.**
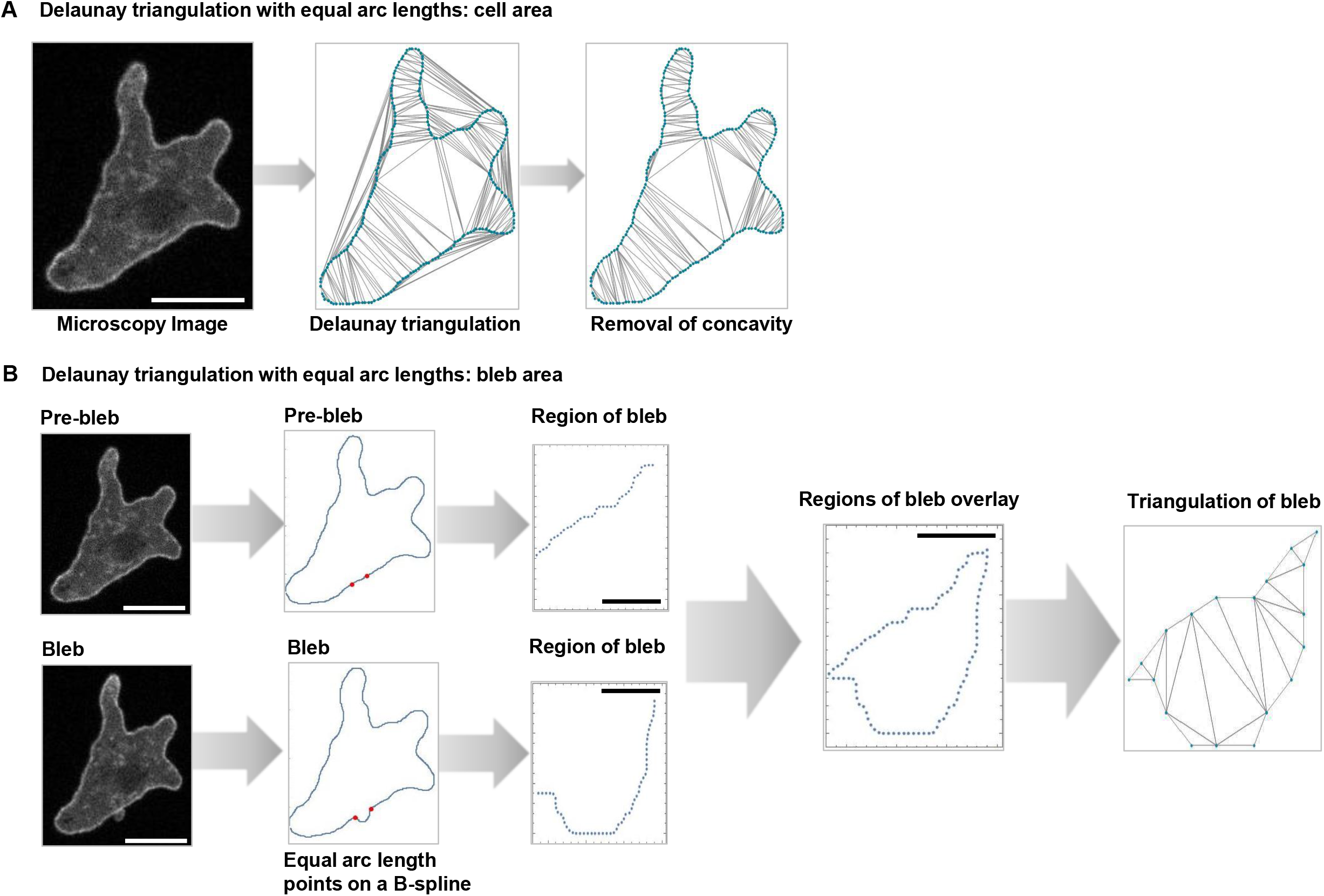
Applying Delauney Triangulation to Determine Cell and Sub cellular Areas. (A) The area of a cell. A cell triangulated by the Delaunay procedure. The convex hull is included within the triangulation. The image on the right shows the result of removing the external triangles. The sum of the triangle areas in this image estimates the cell area. (B) The area of a belb. The two images on the left show the cell before and after the bleb. The next pair (above and below) show the cell by plotting the equi-spaced list of points on the boundary. On the right we see the region of the bleb before and after the event. These two images are overlaid in the lower right. This is a geometric view of the bleb. Finally, we have triangulated the bleb in preparation to estimating its area. Large scale bars identify 10 micron lengths; small scale bars identify 1 micron.

The triangles are either interior or exterior. To remove the exterior ones, we use a procedure based on the Jordan curve theorem. For interior triangles, a line joining the center of the triangle to an interior point of the cell must cross the boundary an even number of times while the opposite is true for exterior triangles. In order to calculate the intersection of a line and the boundary we must have a parameterized representation for the cell boundary. Note that zero is an even number. We show this view in Fig 7A right.

We test each triangle in turn, removing an exterior triangle each time it is encountered. When the exterior triangles are removed, we add the area of the remaining triangles to produce an approximate area of the cell footprint.

Since we have used the sequence of equally spaced points, then the estimated error is easily calculated. We compute the area between the B-spline and segment joining each pair of points in the sequence. (See [25].) We get the error estimate by adding these values.

Alternatively, we can use Green’s theorem applied to the B-spline representation of the cell boundary. In particular, if *∂C* denotes the B-spline boundary, then area is given by

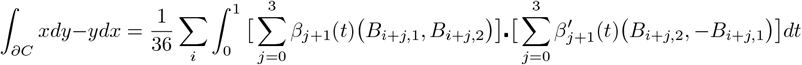

The computed area is the same as with Delauney, only now this approach is less intuitive. However, it is no longer dependent on a built-in function and hence is more easily ported to other programming languages. For instance if we needed to implement this geometric platform into a High Performance Computing environment.

The area of a bleb is related. The bleb as a geometric object is the region between the actin scar and the membrane with newly formed cortex. As stand in for the actin scar, we use the boundary segment associated to the bleb at the prior frame. We get the area by first identifying the points between the base or shoulder points in the after image, in Fig 7B. Using the equally spaced points, we took the union of the points between *a* and *b* in the pre image and *a′* and *b′* in the post image. We show the bleb as represented in this manner. Analogous to the cell, we compute the area by triangulating the region.

Note that the Green’s theorem approach applies equally well to bleb area.

With this basic platform there are many more measurements that can be made. For instance, the bleb reach or farthest extent is an important measurement. To estimate this value, we could look at the circular arcs fitted to the two cell edge segments representing the pre-belb and post-bleb state. These two circular arcs terminate at (nearly) the same locations. The distance between the two arc mid-points is a reasonable stand in for *bleb reach*.

#### 4.3.4 Measuring Curvature

We want a means to measure curvature. Since the cell boundary returned by the microscopy is irregular or bumpy at the micro level, then the computed curvature values will be somewhat chaotic. Even a coherent boundary segment, one that visually appears concave or convex, will contain points in the segment that measure positive or negative depending on the presence of a micro level irregularity. Fig 8A shows the plot of the curvature values for a cell with positive curvature denoted blue and negative curvature denoted red. Even clearly convex or concave segments are not uniformly positive or negative. Furthermore, even then it is unclear what value to return as the curvature for the segment.

**Fig 8.**
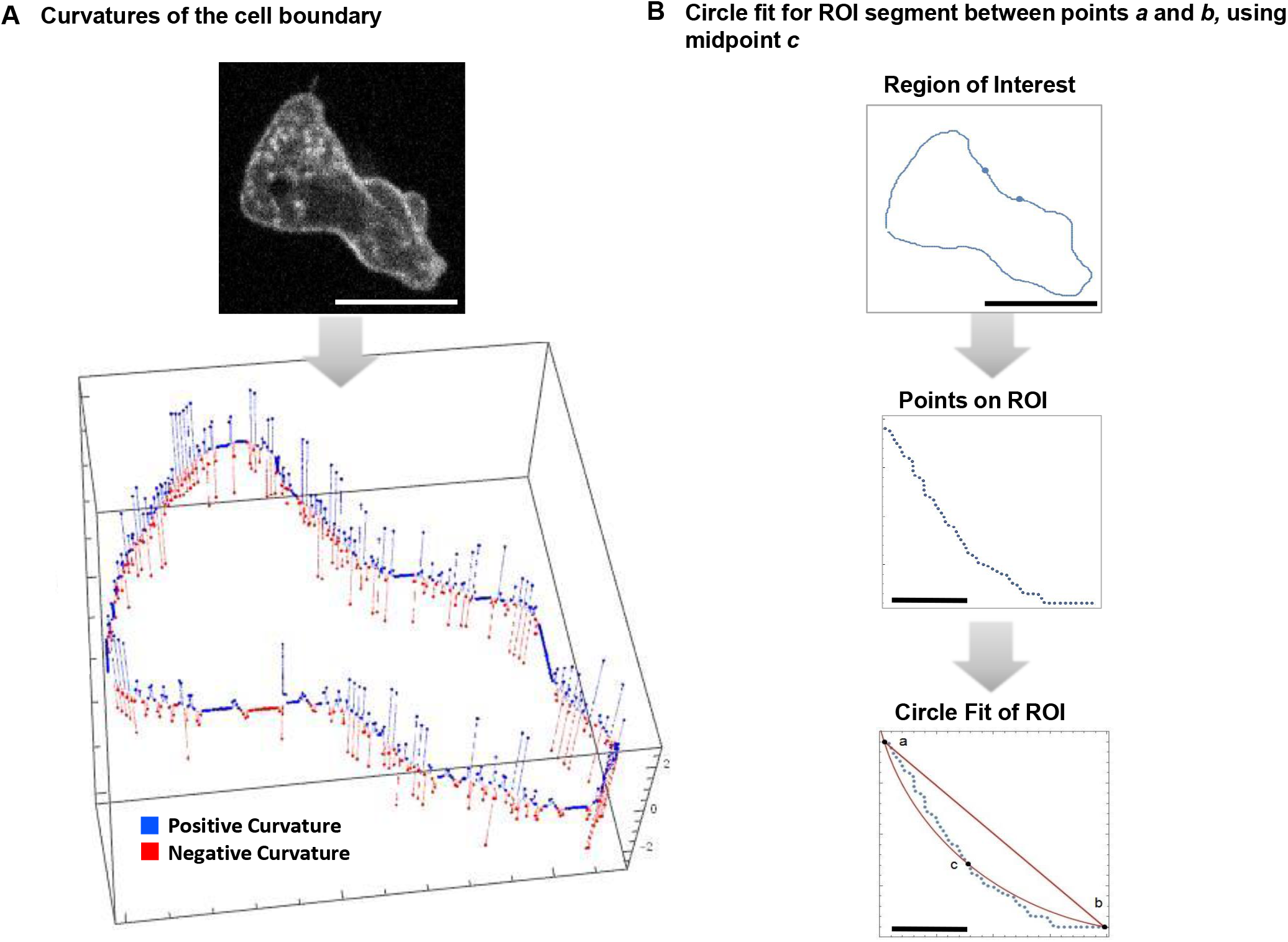
Curvature and Relative Curvature. (A) Cell boundary with positive (blue) and negative (red) curvature values identified. This plot demonstrates the chaotic nature of raw curvature measurements. (B) The relative curvature between the points *a* and *b*. The fitting circle passes through c. The relative curvature value is −0.05097. Scale bars are 10 microns and 1 micron for the ROI.

Another problem with the standard approach to curvature that arcs of the same circle have the same curvature. For *D. discoideum* the length of the circular segment matters. For a given circle a short arc appears to be flat. Such a segment may actually change from concave to convex from frame to frame, whereas a longer segment will remain convex or concave throughout the observational time period.

Our approach to the problem is to fit a circle to the boundary segment. Our procedure is based on the fact that the boundary of the *D. discoideum* cell is formed by internal pressure. In short, it is expected to be locally circular.

Next, we compute the circle arc length between the segment end points, *l_a_*, and compute the linear or Euclidean distance between the segment end points *l_s_*. We use the ratio of those two numbers to measure the segment curvature, *curv_seg_*,

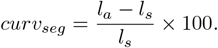

We refer to *curv_seg_* as the *relative local curvature*. In Fig 8A we see that the standard curvature does not identify the segment curvature. Alternatively, relative curvature clearly identifies a concave segment. See Fig 8B. We were successful in reproducing the results of [3].

This does not mean that the computed curvature values are incorrect or useless, only that they are not useful to identify the tendency of segments of the cell boundary.

#### 4.3.5 Classifying blebs by relative local curvature

To illustrate our concept of relative local curvature, we applied it to the problem of classifying blebs according to the curvature of the boundary at the bleb site. We saw results very similar to [3]. In this way, we confirmed our software.

Based on relative local curvature, we classified segments into three categories: Convex, Concave and Low Curvature. Relative local curvature values for concave segments carry a negative sign to distinguish them from convex segments. Since smaller absolute values indicate segments are flatter, a threshold of ±0.005 was used to classify low curvature segments. We used this scheme to categorize segments that bleb in the next frame. We justified the use of a low curvature classification as segments in this range were often slightly convex or concave at one frame and slightly concave or convex at the next. Percentage of blebs occurring in each segment type is given in Table 1, row 1. The total number of cells in this study was 42 with 104 blebs.

In order to get an approximation of the proportion of segments falling under each category, we looked at randomly selected segments from 35 of the cells as shown in Table 1 row 2. Each random segment had the same arc length as a randomly selected bleb. It was selected from a randomly chosen cell/frame.

Table 1 shows the percentage of blebs and random segments belonging to each category. The cells included were wild type and moving under 0.7%agarose. Note that the random segment data reflect the geometry of the cell.

**Table 1.** Bleb distribution, Wild Type and 0.7% Agarose.

| Category | Concave | Low Curvature | Convex | Cells |
| --- | --- | --- | --- | --- |
| Blebs | 51 % | 24.4 % | 24.6 % | 42 |
| Random Segments | 26 % | 32 % | 42 % | 35 |

Statistically, the cutoff of ±0.005 was significant with *p* = 2.913e^−5^. The histogram included in Fig 9A shows clearly that most blebs occur at boundary segments with low negative curvature. The mean relative curvature for concave blebbing segments was –0.09 with standard deviation of 0.11. To intuitively understand these relative curvature values, in Fig 9B we show curves with relative curvature −0.09 and mean plus 1 standard deviation, –0.2.

**Fig 9.**
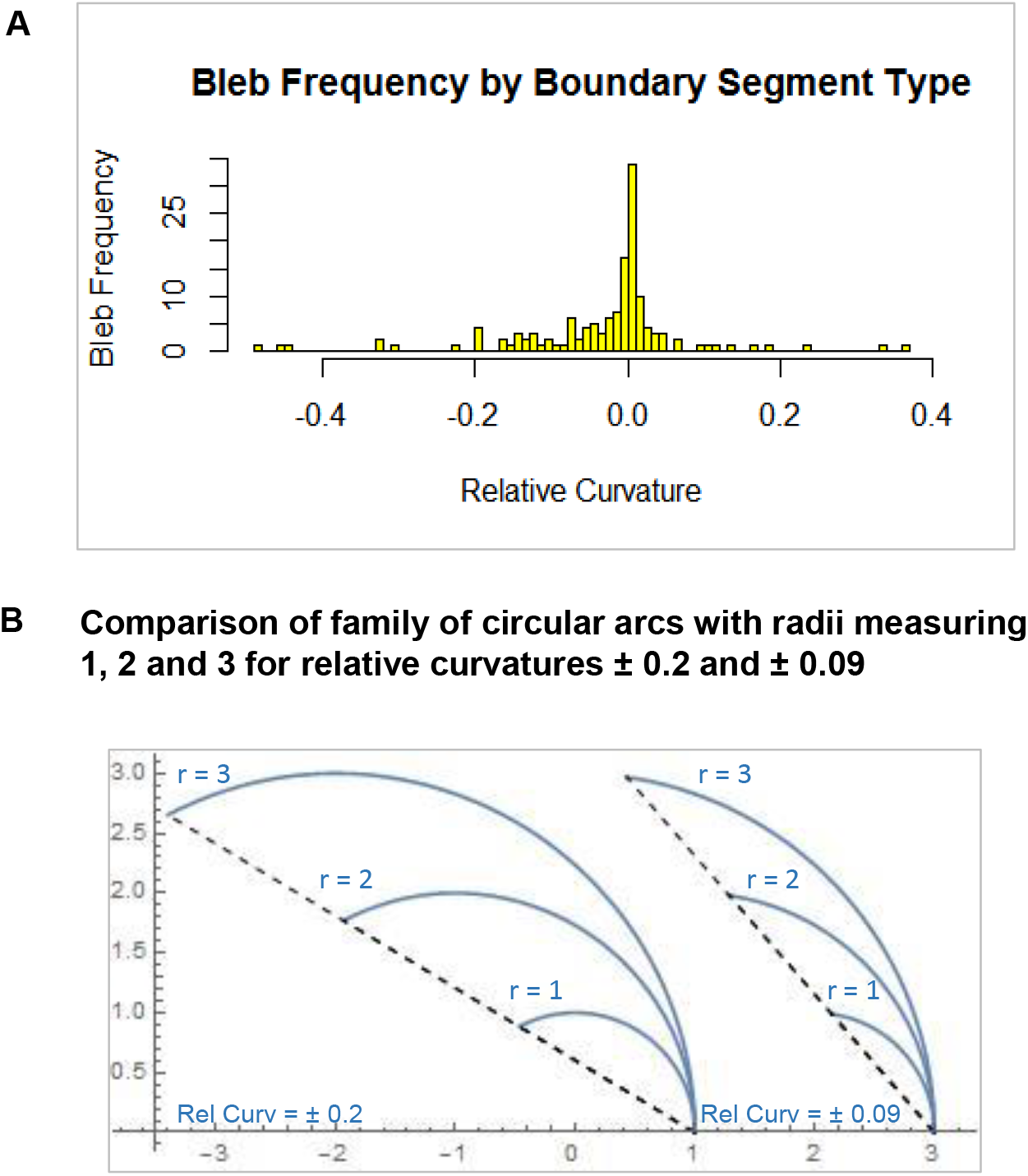
Relative Curvature and Blebbing. (A) Histogram showing the distribution of blebbing sites by local relative curvature. The mean relative curvature for concave blebbing sites is 0.9 with standard deviation 0.11. (B) Examples of relative curvature. The first is 0.09 the mean relative curvature identified in the histogram. The second is 0.2 = 0.09 + 0.11, the mean plus one standard deviation.

The data confirms the work of [3] that blebs are more likely to occur in concave segments even though there are fewer concave segments. As noted in [3], this is a consequence of Young-Laplace equation [26] relating curvature and force. Furthermore, the statistical analysis justifies the use of the three edge segment categories defined by the cutoff ±0.005.

## 5 Discussion/Conclusion

We have advanced current techniques to analyze images of the cell cortex and plasma membrane during blebbing. Through the use of a semi-automated computer application, we have created a firm foundation for our geometric modeling tools. Our technical platform uses accessible software packages ImageJ (https://imagej.nih.gov/ij/) and *Mathematica* (https://www.wolfram.com/mathematica/). Both programs allowed us to answer a variety of biological questions requiring precise measurements of geometric properties of cells. Using *D. discoideum* as our model organism, we were able to monitor and confirm bleb initiation through quantitative measurements of the location of the membrane and the cortex. Using our geometric techniques, we accumulated comprehensive measurements of the curvature of the membrane and cortex throughout bleb formation. From these measurements, we can further investigate bleb characteristics such as bleb area, size, and shape. This provides the tools necessary for further studies, such as cortex tension and membrane energy.

Our methods are not limited to blebbing, but can answer questions regarding cytoskeleton and membrane dynamics in fixed or live cells from actin stains and/or lipid stains. Furthermore, our methods can shed insight into the dynamics of pinocytosis, endocytosis, phagocytosis, and vesicle trafficking by determining the amount of curvature the membrane must endure to initiate these tasks. In the same vein, actin-based structures can be analyzed as well such as the dynamics of the cortex, curvature of filopodia, curvature and area of pseudopodia and lamellipodia, and the shape of endocytic cups. Lastly, our methods facilitate measuring cells on a whole-cell level as well as taking measurements of a region of interest in a cell. Thus, our methods are ideal for those wishing to address biological questions using geometric analysis as these programs are routinely developed from available and accessible software as described. Given that precise measurements and accurate representations of cellular structures are critical first steps for creating realistic mathematical models, our biological/mathematical interface is carefully justified. Hence, we provide fundamental techniques that can improve the quality of many models of various cellular processes.

## Supporting information

SFig 1

SFig 2

SFig 3

SFig 4

SFig 5

## 6 Supplemental Material

### 6.1 Appendix A: Intensity plots of a region without a bleb and full cell videos

SFig 1. ImageJ intensity plots quantitatively confirm membrane and cortex positions of a region without a bleb. Using the same frame of the cell in Fig 2, intensity plots of the signals along the drawn line (white) for each channel show that in a region without a bleb, the membrane and cortex are observed to be in close proximity. Cortex and membrane positions are indicated by (white arrow head) and (black arrow), respectively.

SFig2. ImageJ Intensity plots confirmed the sustained cortex-to-membrane complex in a region without a bleb over time. Using the same frames of the cell in Fig 4, intensity plots were made of a region where there was no bleb (white line). Merges of GFP and RITC time points (*T*_0_ – *T*_3_ for bleb characteristics) with their respective intensity plots (dashed and solid lines) confirmed a sustained cortex-to-membrane complex over time. Cortex and membrane positions are indicated by (white arrow head) and (black arrow), respectively.

SFig 3. A bleb was identified through actin debris under 0.5 percent agarose. Top row shows the frame before the bleb, and the bottom row shows the frame with the bleb in progress, identified by the actin debris present in the bleb. Left, in the GFP channel, the white arrow indicates the region where the actin debris will be present in both time frames. Center, the RITC channel shows membrane position in both frames with the bleb region present in the second time frame. Right, the merge between both GFP and RITC channels false colored green and red, respectively. Images were taken on the 100x objective.

SFig 4. A bleb was identified through actin debris under 0.7 percent agarose. Top row shows the frame before the bleb, and the bottom row shows the frame with the bleb in progress, identified by the actin debris present in the bleb. Left, in the GFP channel, the white arrow indicates the region where the actin debris will be present in both time frames. Center, the RITC channel shows membrane position in both frames with the bleb region present in the second time frame. Right, the merge between both GFP and RITC channels false colored green and red, respectively. Images were taken on the 80x objective.

SFig 5. A bleb was identified through actin debris under 0.7 percent agarose without RITC-Dextran. Left, the frame before the bleb. Center, the frame with the bleb in progress, identified by the actin debris present in the bleb. Right, the next frame showing the cortex reforming in the bleb. Images were taken on the 100x objective.

Video 1. Microscopy video of Figs 2–4’s cell. This wild-type cell expressing Life-Act-GFP crawled upward a cAMP gradient under a 0.5 percent agarose gel laced with RITC-Dextran. The exposure of the GFP and RITC channels were 0.800 and 0.122 secs, respectively. However, due to the microscope’s limitations in the speed of changing channels, the lowest frame rate achievable was 1.66 secs.

Video 2. Microscopy video of Fig. 5’s cell. This wild-type cell expressing LifeAct-GFP crawled upward a cAMP gradient under a 0.7 percent agarose gel without RITC-Dextran to visualize bleb progress after nucleation. With this assay, the frame rate for this was 0.800 secs. This lowered frame rate allowed for observing bleb progression over time via the cytoplasm with labeled via residual F-actin LifeAct-GFP labeled debris filling the bleb.

Video 3. Microscopy video of Fig. 6A and Fig. 8’s cell. Another wild-type cell expressing LifeAct-GFP crawled upward a cAMP gradient under a 0.7 percent agarose gel without RITC-Dextran. The framerate was 0.800 secs.

Video 4. Microscopy video of Fig.6B and Fig. 7’s cell. Another wild-type cell expressing LifeAct-GFP crawled upward a cAMP gradient under a 0.7 percent agarose gel without RITC-Dextran. The framerate was 0.800 secs.

### 6.2 Appendix B: Convergence Result for B-Splines

See [25] for a development of B-Splines. The following argument is outlined in that reference.

Let *f* be a function defined on an interval [*a, b*]. *f* is Lipschitz continuous provided there is a constant *K* with |*f*(*c*) – *f*(*d*)| < *K*|*c – d*| for any *c,d* ∈ [*a, b*]. We want to prove the following statement:

*Let f be Lipschitz continuous, real valued function on an interval*, [*a, b*] *and let a* = *α*_0_ < *α*_1_ < … < *α_n_* = *b be a partition of the interval with mesh parameter, h* = max_*i*_(*α_i_* – *α*_*i*–1_). *Suppose that σ is a B-spline supported by the point set* (*α_i_, f*(*α_i_*)), *then σ converges to f in L*^∞^ *norm as h* → 0. *In particular, if* (*x, f*(*x*)) *is a point on the graph of f and* (*x, y*) *is the corresponding point on σ, then* ║(*x, f*(*x*)) – (*x,y*)║ < 12*Kh, where K is the Lipschitz constant*.

Let *β_i_*(*t*) be a B-spline basis function (*i* = 1, 2, 3, 4) and set *L* = max_*i,t*_ |*β_i_*(*t*)|, for *t* in [0,1]. It is routine to verify that L < 1.

We begin convergence. We take *x* in [*a, b*] and suppose that *x* is the first coordinate of a point on the B-spline. In particular, there is a *k* so that *x* associates to the *k^th^* B-spline segment, *σ_k_*. Hence, there is a *t*_0_ in the unit interval with the first coordinate of *σ_k_*(*t*_0_) equal to *x*. Let *y* denote the second coordinate. In addition, if *γ* is the first coordinate of *σ_k_*(0) and *δ* is the corresponding coordinate of *σ_k_*(1), then

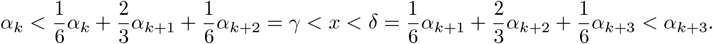

Therefore, |*x* – *α*_*k*+*j*_| ≤ 3*h* for *j* = 0, 1, 2, 3.

We verify ║(*x, f*(*x*)) – (*x,y*)║ < 12*KLh* in Euclidean norm. Toward this end, we calculate,

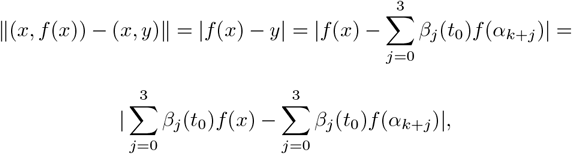

since 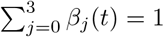. Continuing,

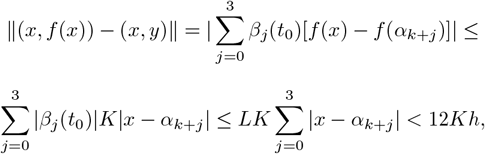

since *x* ∈ [*α_k_, α*_*k*+3_]. The proposition now follows.

## Acknowledgments

We acknowledge Dr. Emmanuel Asante-Asamani, Department of Biological Sciences and Department of Mathematics and Statistics, Hunter College, CUNY, for his contribution to the LaTex typesetting and the mathematical content. In addition, we acknowledge the contribution of L. Leslie Liu, Department of Mathematics and Statistics, Hunter College, CUNY, for assistance with the statistical analysis.

This work was supported by grants to Derrick Brazill from the National Science Foundation (MCB-1244162), a PSC-CUNY grant (69271 00 47), as well as Research Centers in Minority Institutions Program grants from the National Institute on Minority Health and Health Disparities (8 G12 MD007599) from the National Institutes of Health.

## References

1. K. Yoshida and T. Soldati, Dissection of amoeboid movement into two mechanically distinct modes, J. Cell Sci., 2006;119, pp. 3833–3844.

2. T. Lammermann, and M. Sixt, Mechanical nodes of ‘amoeboid’ cell migration, Curr. Op. Cell Bio., 2009;21, pp. 636–644.

3. R. Tyson, E. Zatülobskiy, R. Kay, and T. Bretschneider, How blebs and pseudopods cooperate during chemotaxis, PNAS, 2014;32, pp. 11703–11708.

4. A. Otto, H. Collins-Hooper, H. Patel, P. Dash, and K. Patel, Adult skeletal muscle stem cell migration is mediated by a blebbing/amoeboid mechanism, Rejuvenation Res., 2011;14, pp. 249–260.

5. A. Diz-Münoz, P. Romanczük, W. Yü, M. Bergert, K. Ivanovitch, G. SalbreüX, C. Heisenberg, and E. PalüCH, Steering cell migration by alternating blebs and actinrich protrusions, BMC Biology, 2016;14, p. 74.

6. H. Blaser, M. Reiohman-Fried, i. Castanon, K. Ðümstrei, F. Marlow, K. Kawakami, L. Solnioa-Krezel, and C. Heisenberg, Migration of zebrafish primordial germ cells: A role for myosin contraction and cytoplasmic flow, Development Cell, 2006;160, pp. 613–627.

7. K. Wolf, I. Mazo, H. Leüng, K. Engelke, U. Von Andrian, E. Deryugina, a. s. E. Brooker, and P. Friedl, Compensation mechanism in tumor cell migration: mesenchymal-amoeboid transition after blocking of pericellular proteolysishe, J, Cell Biol., 2003;160, pp. 267–277.

8. H. Yamaguohi and J. Condeelis, Regulation of the actin cytoskeleton in cancer cell migration and invasion, Biochim Biophys Acta, 2007;1773, pp. 642–652.

9. B. Maügis, J. Brügües, P. Nassoy, N. G. P. Sens, and F. Amblard, Dynamic instability of the intracellular pressure drives bleb-based motility, J. Cell Sci., 2010;123, p. 18.

10. J. Nichols, Ð. Veltman, and R. Kay, Chemotaxis of a model organsim: progress with dictyostelium, Curr. Op. Cell Bio., 2015;36, pp. 7–12.

11. G. Charras, M. Coughlin, T. Mitohison, and L. Mahadevan, Life and times of a bleb, Biophys. J., 2008;95, pp. 1836–1853.

12. T. Wooley, E. Gaffney, J. Oliver, S. Waters, and R. Baker, Global construction or local growth, bleb shape depends on more than just cell structure, J. Theoretical Bio., 380 (2015), pp. 83–97.

13. R. Guy and W. Stryohalsky, Intracellular pressure dynamics in blebbing cells, Biophys. J., 2010;110, pp. 1168–1179.

14. W. Stryohalsky, C. Copos, O. Lewis, and R. Guy, A proelastic immersed boundary method with applications to cell biology, J. Comp. Phys., 2015;282, pp. 77–97.

15. S. Collier, P. Pashke, R. Kay, and T. Bretsohneider, Image based modeling of bleb site selection Sci. Rpts, 2017; DOI: 10.1038.

16. I. Markela, V. Srivastava, Ð. Robinson, and R. Gagnon, Cell Blebbing in confined microfludic Environments, PLOS One, 2015;DOI: 10.1371.

17. E. Paluoh and E. Raz, The role of regulation of blebs in cell migration, Curr Op in Cell Bio, 2013;25, pp. 582–590.

18. E. Zatülovskiy, R. Tyson, T. Bretsohneider, and R. Kay, Bleb-driven chemotaxis of Dictyostelium cells, J. Cell Biol., 2014;204, pp. 1027–1044.

19. C. Choi, H. Patel, and Ð. Barber, Expression of Actin-interacting Protein 1 Suppresses Impaired Chemotaxis of Dictyostelium Cells Lacking the Na+-H+ Exchanger NHE1, Mol. Biol. Cell., 2010;21, pp. 162–170.

20. J. Riedl, A. Crevenna, K. Kessenbrook, J. Yu, Ð. Neükirohen, M. Bista, F. Bradke, Ð. Jenne, T. Holak, Z. Werb, M. Sixth, and R. Wedlioh-Soldner, Lifeact: a versatile marker to visualize F-actin, Nat. Methods, 2008;5(7), pp. 605–607.

21. M. Pang, a. Lynes, and Ð.A. Kneoht, Variables controlling the expression level of exogenous genes in Dictyostelium, Plasmid, 2009;41, pp. 187–97.

22. M.G. Lemieüx, D. Janzen, R. Hwang, J. Roldan, I. Jarchüm, and D.A. Knecht Visualization of the actin cytoskeleton: different F-actin-binding probes tell different stories, Cytoskeleton (Hoboken), 2014;71(3), pp. 157–69, doi: 10.1002/cm.21160.

23. M. Süssman, Cultivation and synchronous morphogenesis of Dictyostelium under controlled experimental conditions, Methods. Cell. Biol., 1987;28, pp. 9–29.

24. I.e. Catrina, S.A. Marras, and D.P; Bratü, Tiny molecular beacons: LNA/2'-O-methyl RNA chimeric probes for imaging dynamic mRNA processes in living cells, ACS Chem Biol., 2012;21(9), pp. 1586–95.

25. J. LoüSTAü, Elements of Numerical Analysis with Mathematica, World Scientific Press, 2017.

26. L. Landaü and E. Lifschitz, Fluid Mechanics, Butterworth and Heineman, 2nd ed., 1987.

