## Supplementary figures and images for "Advances in Geometric Techniques for Analyzing Blebbing in Chemotaxing *Dictyostelium* Cells"

### SFig 1

#### SFig 1. of Fig 3: IP of No bleb region. 8/29/18

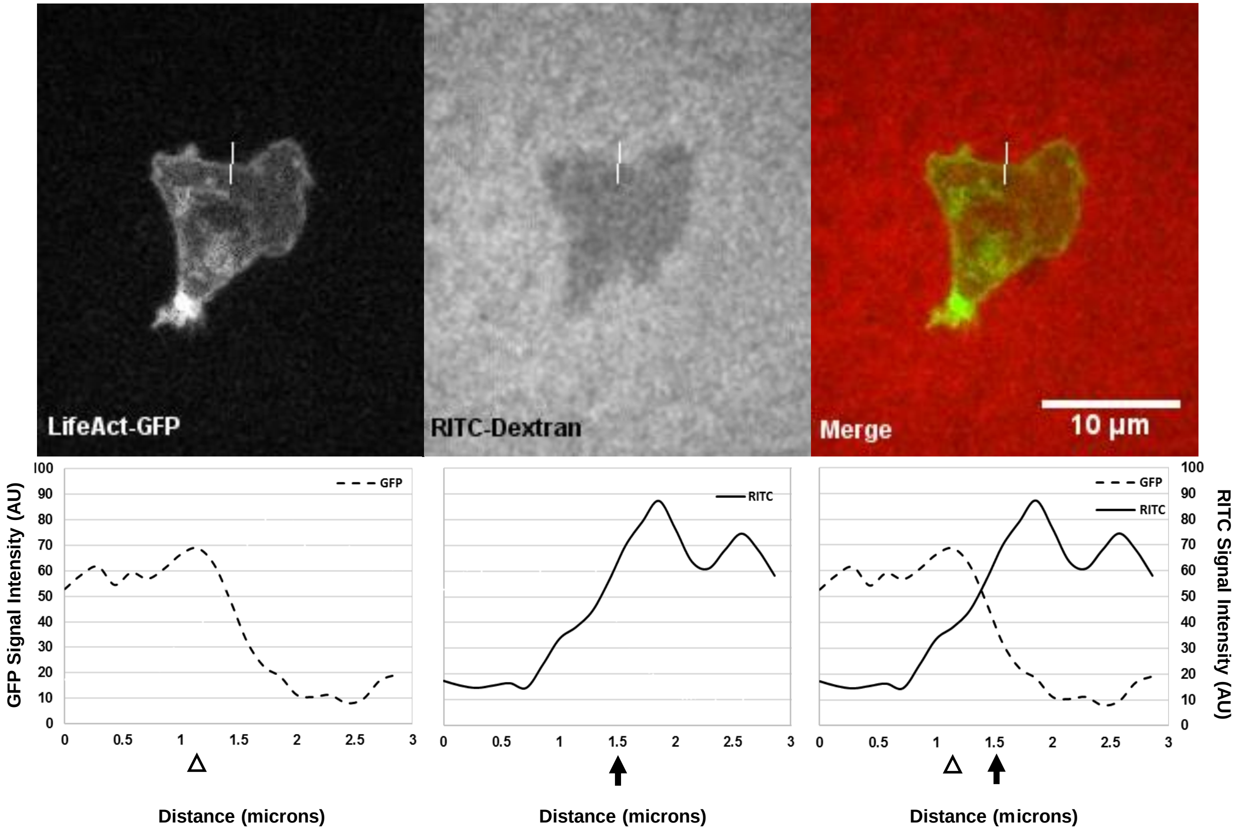

### SFig 2

#### SFig 2. of Fig 4:IP of No bleb region. 8/29/18

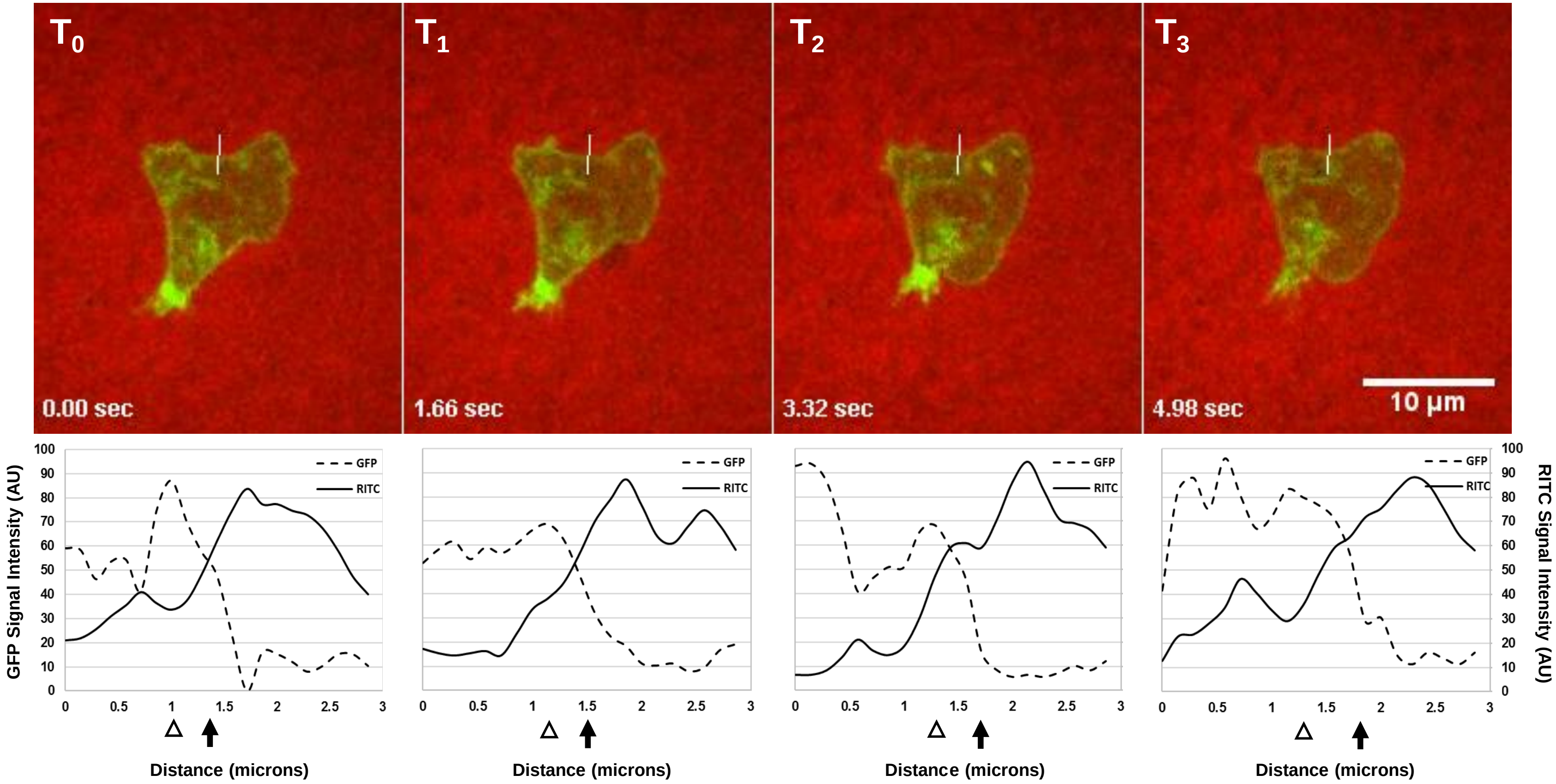

### SFig 3

#### SFig 3. Bleb identification via actin debris under 0.5% agarose assay 100x

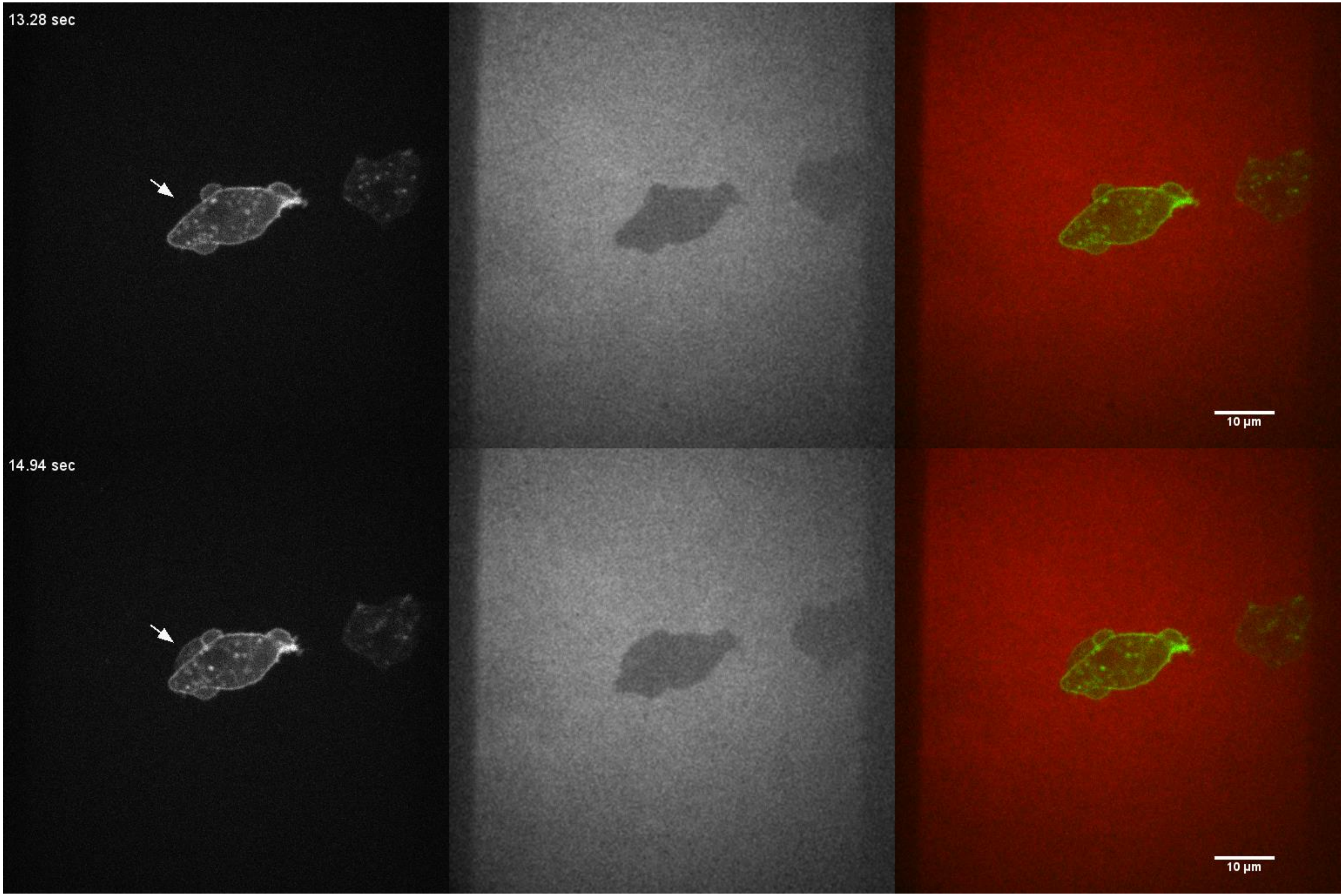

### SFig 4

#### SFig 4. Bleb identification via actin debris under 0.7% agarose assay 80x

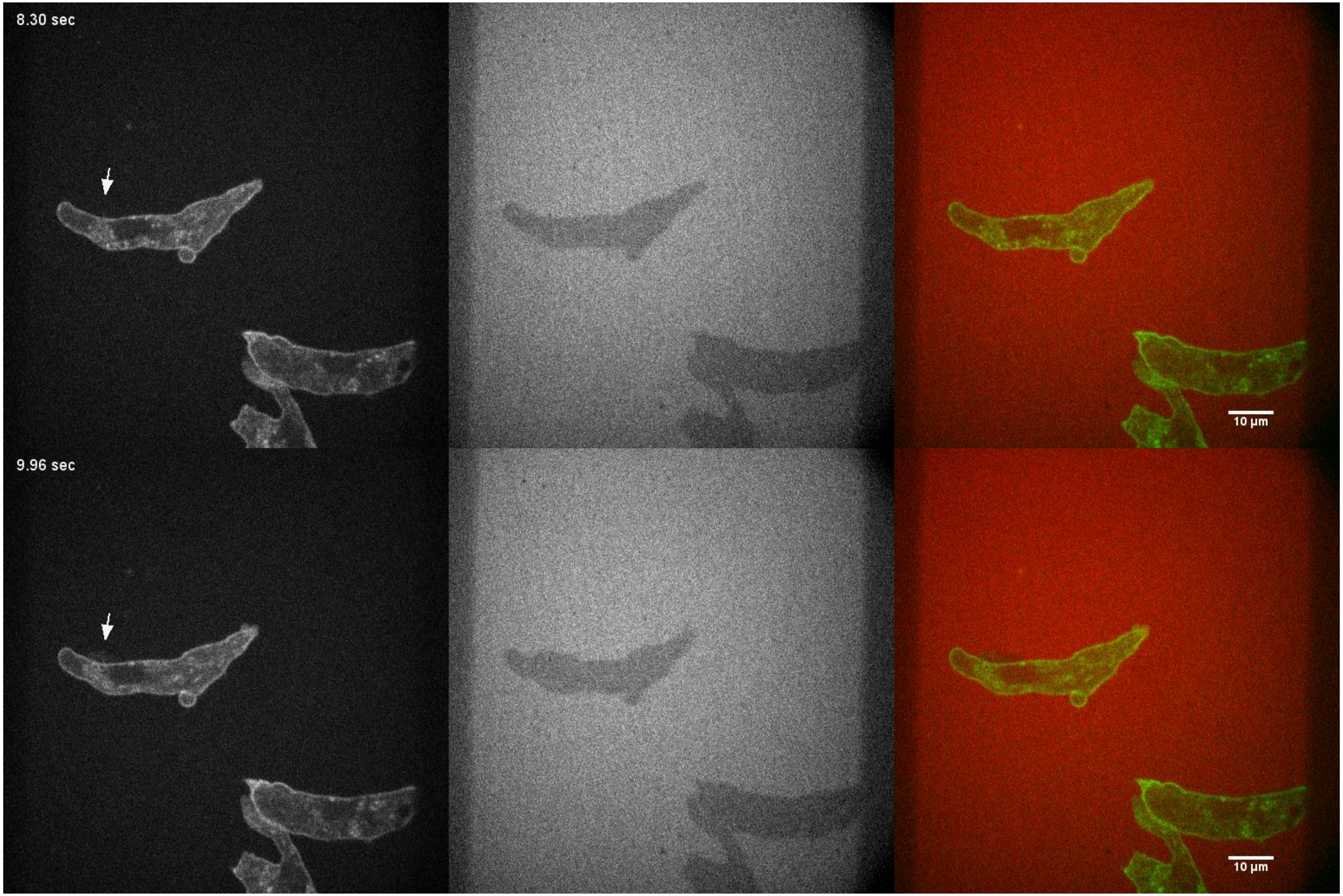

### SFig 5

#### SFig 5. Bleb identification via actin debris under 0.7% Flow 100x

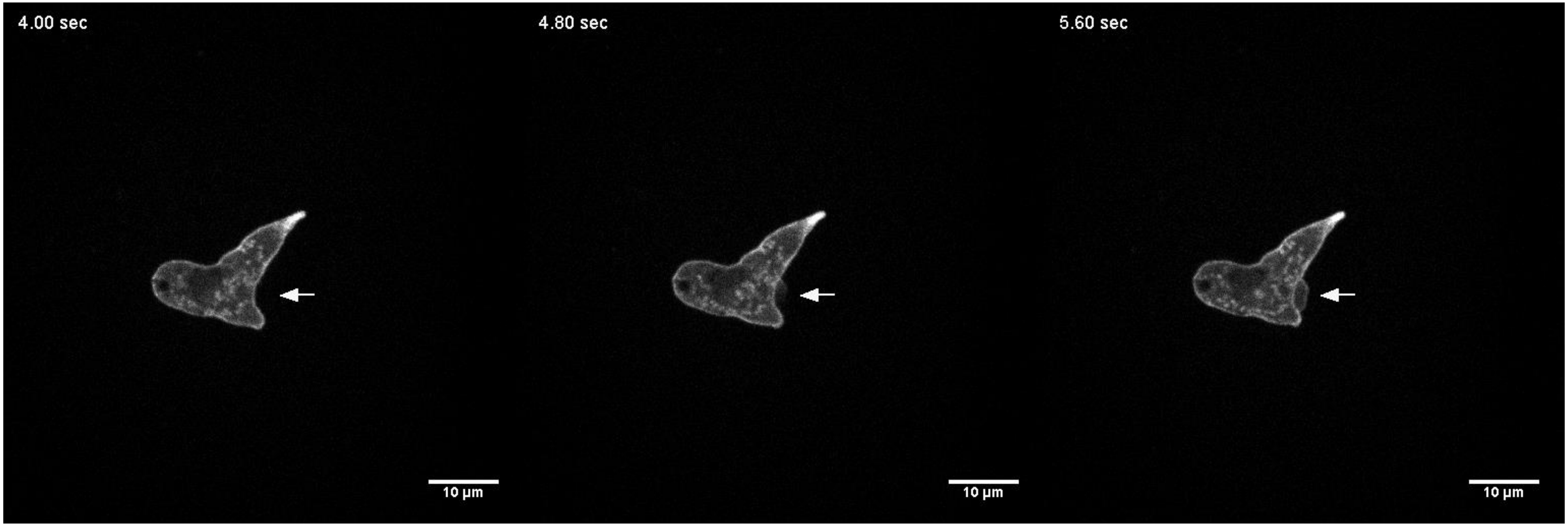
